# Mitogen-Activated Protein Kinase-Dependent Fiber-Type Regulation in Skeletal Muscle

**DOI:** 10.1101/361600

**Authors:** Justin G. Boyer, Taejeong Song, Donghoon Lee, Xing Fu, Sakthivel Sadayappan, Jeffery D. Molkentin

## Abstract

Mitogen-activated protein kinases (MAPK) are conserved protein kinases that regulate a diverse array of cellular activities. Stress or mitogenic signals activate three primary branches of the greater MAPK cascade, each of which consists of a phosphorylation-dependent array of successively acting kinases. The extracellular signal-regulated kinase 1/2 (ERK1/2) branch is regulated by growth factory signaling at the cell membrane, leading to phosphorylation of the dual-specificity kinase MEK1, which is dedicated to ERK1/2 phosphorylation. Previous studies have established a link between MAPK activation and endurance exercise, but whether a single MAPK is responsible for establishing muscle metabolic fate is unclear. Using mouse genetics we observed that muscle-specific expression of a constitutively active MEK1 promotes greater ERK1/2 signaling that mediates fiber-type switching in mouse skeletal muscle to a slow, oxidative phenotype with type I myosin heavy chain expression. Induced expression of the activated MEK1 mutant using either a MyoD-Cre or myosin light chain-Cre strategy equally increased the number of type I fibers in skeletal muscle with significantly reduced size compared to controls. Moreover, activation of MEK1 in mature myofibers of an adult mouse using a transgene containing a tamoxifen inducible MerCreMer cDNA under the control of a skeletal α-actin promoter produced a similar phenotype of switching towards a slow-oxidative program. Physiologic assessment of mice with greater skeletal muscle slow-oxidative fibers showed enhanced metabolic activity and oxygen consumption with greater fatigue resistance of individual muscles. In summary, these results show that sustained MEK1-ERK1/2 activity in skeletal muscle produces a fast-to-slow fiber-type switch, suggesting that modulation of this signaling pathway may represent a therapeutic approach to enhance the long-term metabolic effectiveness of muscle *in vivo*.

## Introduction

Skeletal muscles vary on the basis of their shape, size and physiologic characteristics that collectively determine their function. Myofibers are individual contractile units that compromise all muscles and they are grossly categorized as either type I slow-twitch oxidative, type IIA fast-twitch oxidative-glycolytic myofibers and type IIB/IIX fast-twitch glycolytic myofibers [1]. These fiber types have distinct molecular and functional properties and can be identified at the histological level by expression of specific myosin heavy chain (MyHC) isoforms and selected metabolic genes. For example, as an adaptation to continual usage, type I fibers express slow-twitch-specific contractile proteins such as *Myhc7* gene product and are rich in myoglobin and mitochondria to produce a fatigue resistance profile compared with fast-twitch fibers that are more specialized for bursts of activity [1]. The desire to better understand the molecular mechanisms regulating a fast-to-slow myofiber switch has been fueled by the potential therapeutic value of a more oxidative, metabolic state in chronic disease such as obesity and type 2 diabetes [2]. Type I fibers also appear to better protect skeletal muscle from genetic diseases such as Duchenne muscular dystrophy [3, 4]. Potent regulators of the slow-oxidative program include the calcineurin-signaling pathway [5, 6], the AMP-activated protein kinase (AMPK) [7], the peroxisome proliferator-activated receptor β/δ pathway [8], as well as the transcriptional co-activator PGC-1α pathway [9].

Mitogen-activated protein kinases (MAPK) are a highly conserved network that transduce extracellular signals into an intracellular responses involving three to four tiers of kinases that constitute specific amplifying phosphorylation cascades. The cascade culminates in the phosphorylation and activation of effector kinases, p38, c-Jun N-terminal kinases (JNK1/2) and extracellular signal-regulated kinases 1/2 ERK1/2 [10]. In the case of ERK1/2 activation, the rapid accelerated fibrosarcoma (Raf) kinase is recruited and activated by the GTPase RAS at the cell membrane leading to linear phosphorylation events in which Raf activates MEK1/2 kinases that then directly phosphorylate ERK1/2 [11, 12]. Although the unique substrate specificity of the Raf-MEK1/2-ERK1/2 pathway has been previously demonstrated, ERK1/2 can, upon activation, translocate to the nucleus and phosphorylate a large number of substrates [11]. ERK1/2 have conventionally been associated with regulating cell proliferation and cell survival [12], however, in post-mitotic differentiated cells the role of ERK1/2 can vary. In cardiomyocytes for example, ERK1/2 signaling regulates eccentric versus concentric cardiac growth [13, 14].

In skeletal muscle, a strong correlation between exercise and ERK1/2 activation exists. For example, in a study involving human subjects, Aronson et al. reported that participants using a one-legged exercise protocol had increased ERK1/2 activation in the exercised leg relative to the contralateral leg [15]. Furthermore, research on ERK1/2 in skeletal muscle revealed an increase in phosphorylated ERK1/2 in marathon runners [16]. With respect to ERK1/2 manipulation *in vivo*, Shi et al. transfected a plasmid expressing MAPK phosphatase-1 (MKP-1) into the gastrocnemius of mice, which showed an increase in type I fibers, suggesting that inhibition of ERK1/2 causes a fast-to-slow fiber-type conversion [17]. However, MKP-1 is not specific to ERK1/2 as it also inhibits JNK1/2 and p38 when overexpressed *in vivo* [18]. Activation of both p38 and JNK MAPKs are also observed following bouts of acute exercise or marathon running, suggesting that these kinases could play a role as well [19]. Indeed, Pogozelski et al. showed that muscle-specific deletion of p38γ in mice caused a reduction in exercise-induced oxidative fiber-type transition [20]. In the present study, we have taken advantage of mouse genetics to reveal that MEK1, which exclusively phosphorylates and activates ERK1/2, leads to a robust induction of the type I, oxidative muscle phenotype in mice.

## Results

### MEK1-ERK1/2 signaling-dependent fast-to-slow fiber-type switch

To determine if the activation of MEK1-ERK1/2 signaling regulated the fiber-type program in skeletal muscle we utilized a mouse containing a Cre-dependent constitutively active MEK1 cDNA (*Map2k1* gene product) inserted into the *Rosa26* locus (Fig.1A). *Rosa26*-MEK1 mice were crossed with *Myl1*-Cre gene-targeted mice (referred to as MLC-Cre) which expresses Cre-recombinase in differentiating myofibers [21]. MEK1 expression levels and an increase in total ERK1/2 were observed in various hindlimb skeletal muscles (Fig 1B, Supplementary Material, Fig. S1A-D). The increase in total ERK1/2 protein observed in the *Rosa26*-MEK1 × MLC-Cre skeletal muscle-specific mice was previously observed in cardiac-specific MEK1 transgenic mice [22].

**Figure 1.**
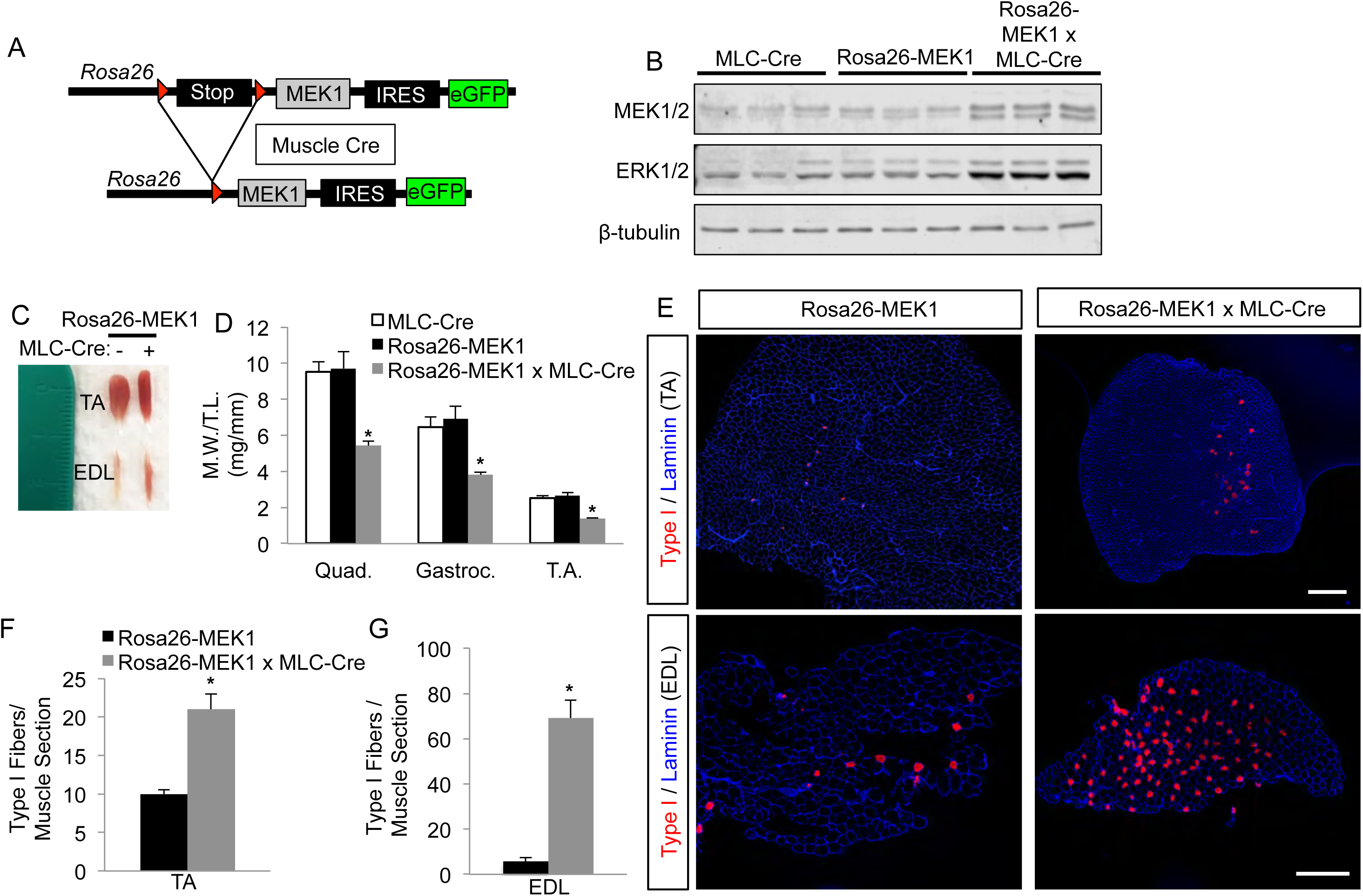
Constitutive active MEK1 expression leads to an increase in type I fibers. (A) Schematic representation of the constitutively active MEK1 cDNA targeted to the *Rosa26* locus that has flanking loxP sites (red) and is only expressed after Cre-recombinase dependent recombination. (B) Western blot analysis for the indicated proteins using gastrocnemius lysate from 6 month-old mice (n = 3). (C) Representative image of the tibialis anterior (TA) and extensor digitorum longus (EDL) muscles from 6 month-old *Rosa26*-MEK1 and *Rosa26*-MEK1 × MLC-Cre animals. (D) Relative muscle weights (M.W.) of quadriceps (Quad), gastrocnemius (Gastroc.) and TA muscles normalized to tibia length (T.L.) from 6 month-old control and *Rosa26*-MEK1 × MLC-Cre mice. n = 4-6 mice per group. ^*^P < 0.05 versus controls. (E) Representative histological sections from TA and EDL muscles immunostained for the MYH7 protein (red) to identify type I fibers and for laminin (blue) to delineate myofiber outlines in histological sections from mice of the indicated genotypes. Scale bars: 500 μm. (F-G) Quantification of slow-oxidative type I fibers across an entire muscle histological section at the mid-belly from the TA (F) and the EDL (G) taken from 6 month-old mice of indicated genotypes. n = 3-6 mice per group. ^*^P < 0.05.

Remarkably, skeletal muscle from *Rosa26*-MEK1 × MLC-Cre mice showed a prominent increase in red color compared with Rosa26-MEK1 only controls, suggesting a slow oxidative program switch (Fig 1C). This difference was evident at 2 months of age (Supplementary Material, Fig. S2A) and 6 months of age (Fig. 1C). Relative muscle weights from the quadriceps, gastrocnemius and tibialis anterior (TA) muscles were also significantly reduced in *Rosa26*-MEK1 × MLC-Cre animals compared to controls at 2 months of age (Supplementary Material, Fig. S2B). At 6 months of age the differences in size was even more pronounced (Fig. 1D).

Muscle histological sections collected and stained for type I myosin heavy chain (*Myh7* gene product) to identify fibers that were associated with the slow-twitch program in the soleus (Supplementary Material, Fig. S2C), TA (Fig. 1E) and the extensor digitorum longus (EDL, Fig. 1E and Supplementary Material, Fig. S2C). We also quantified the number of type I positive fibers per muscle section. Compared to *Rosa26*-MEK1 controls, we observed a significant increase in type I fibers in 3 muscles tested: the TA (Fig. 1F and Supplementary Material, Fig. S2D), the soleus muscle (Supplementary Material, Fig. S2E) and the EDL muscle (Fig. 1G and Supplementary Material, Fig. S2F). These results suggest that MEK1-ERK1/2 signaling promotes a slow-oxidative fiber-type program in skeletal muscle.

### ERK2 underlies the slow-twitch phenotype observed downstream of MEK1

We have previously observed a greater need for ERK2 protein (*Mapk1* gene) relative to ERK1 protein (*Mapk3* protein) in the heart in programming adaptive responsiveness [23]. Here we evaluated if ERK1 or ERK2 preferentially contributed to the fast-to-slow fiber-type switch observed in the activated MEK1 expressing mice. We crossed *Rosa26*-MEK1 and MLC-Cre alleles into the *Mapk1* floxed (f) targeted background or separately into the germline *Mapk3*^-/-^ background. For the first cross the MLC-Cre produces recombination of both the *Rosa26*-MEK1 and *Mapk1^f/f^* alleles in myofibers. These mice showed increased MEK1 expression and reduced ERK2 protein (lower band) with a mild increase in ERK1 protein (upper band, Fig 2A). Skeletal muscle weights in the quadriceps, gastrocnemius and TA were unchanged between normal controls and MEK1 expressing mice lacking *Mapk1* (Fig. 2B). More importantly, loss of *Mapk1* now eliminated the ability of MEK1 to augment type I myofiber content (MYH7 protein) compared with controls (Fig 2C,D). By comparison, loss of *Mapk3* gave a compensatory increase in ERK2 protein expression in the presence of the activated *Rosa26*-MEK1 expressing allele (Fig. 2E), and it permitted a decrease in skeletal muscle weights of the quadriceps, gastrocnemius and TA (Fig 2F), as observed in activated MEK1 expressing mice replete with ERK1/2 (Fig 1D). More importantly, loss of *Mapk3* in muscle still produced the large increase in type I fibers (Fig 2G,H). These results suggest that ERK2 is the primary downstream MEK1 kinase responsible for the MEK1-mediated fiber-type switching in mice.

**Figure 2.**
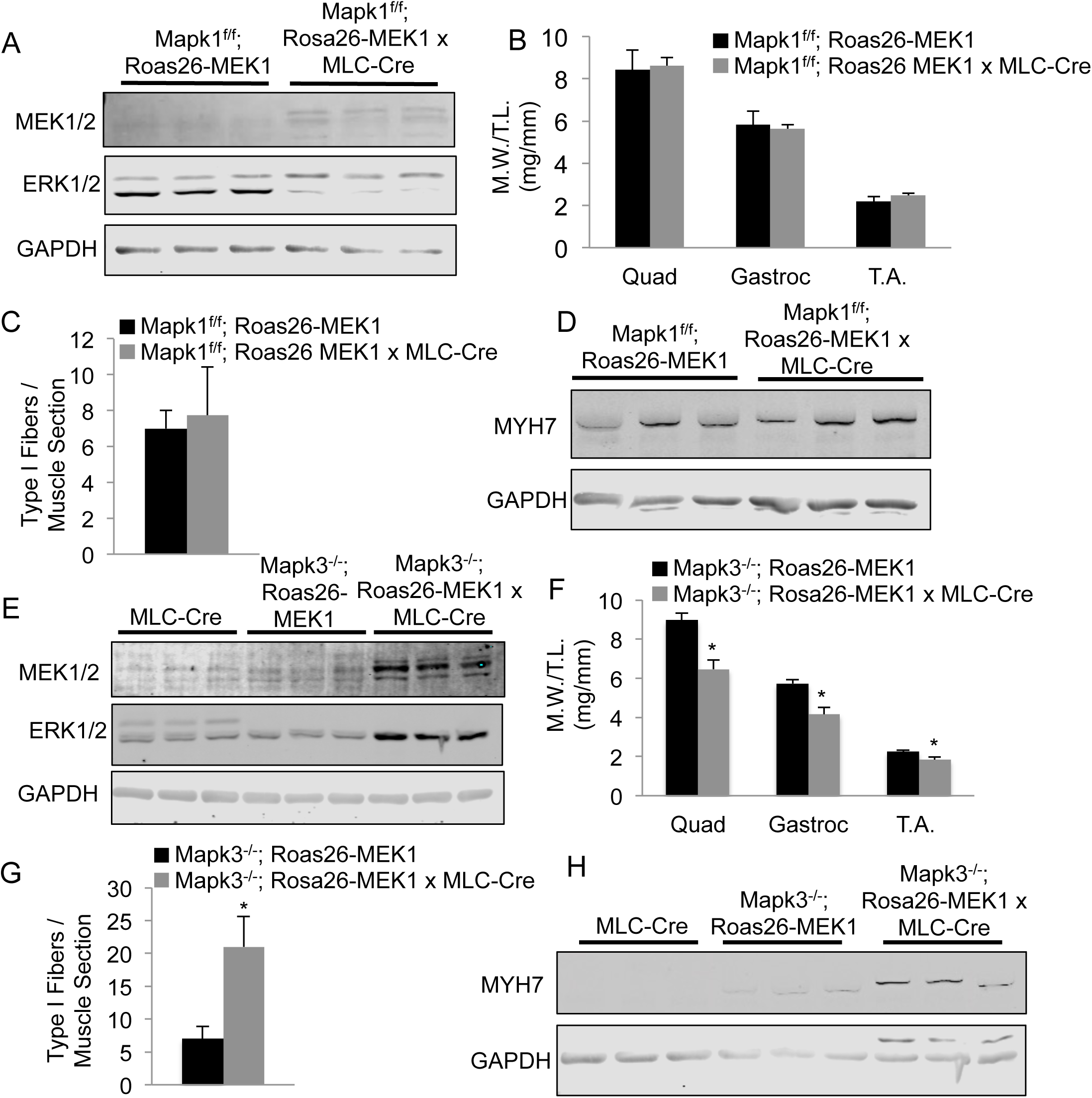
ERK2 drives the MEK1-mediated fast-to-slow fiber-type switch. (A) Western blot analysis for total MEK1/2 and total ERK1/2 protein using lysate from the gastrocnemius muscle from mice of the indicated genotypes (n = 3). GAPDH was used a loading control. (B) Muscle-weight normalized to tibia-length (M.W./T.L.) at 5 months of age for the quadriceps (Quad), gastrocnemius (Gastroc) and TA muscle from mice of the indicated genotypes. n = 5-6 mice per group. (C) Quantification of total type I fibers in an EDL muscle histological section taken at the mid-belly of the muscle from mice of the indicated genotypes. (D) Western blot analysis for MYH7 (type I) using lysate from the gastrocnemius muscle from 5 month-old animals of the indicated genotypes of mice. GAPDH was used a loading control. (E) Western blot analysis for total MEK1/2 and total ERK1/2 using lysate from the gastrocnemius muscle from mice of the indicated genotypes. GAPDH was used a loading control. (F) M.W./T.L. at 5 months of age of the Quad, gastroc, and TA muscle from mice of indicated genotypes. n = 6-7 mice per group. ^*^P < 0.05. (G) Quantification of total type I fibers in an EDL histological section at the mid-belly of the muscle from mice of the indicated genotypes. ^*^P < 0.05. (H) Western blot analysis for MYH7 (type I) using lysate from the gastrocnemius muscle from 5 month-old mice of indicated genotypes (n = 3). GAPDH was used a loading control.

### MEK1 drives the slow oxidative program in myoblast progenitors and adult muscle

We crossed the *Rosa26*-MEK1 allele-containing mice with mice containing the MyoD^iCre^ targeted insertion allele [24]. This strategy allowed us to induce MEK1-ERK1/2 signaling at an early stage in muscle progenitors of the developing embryo, irrespective of initial fiber-type [24]. Skeletal muscles from *Rosa26*-MEK1 × MyoD-Cre mice still showed a noticeably more oxidative red appearance compared with *Rosa26*-MEK1 only littermate controls (Fig. 3A). At 6 months of age, relative muscle weights were also significantly decreased in Rosa26-MEK1 × MyoD-Cre mice compared to controls (Fig. 3B), similar to the effect observed with the MLC-Cre allele (Fig 1D). At the histological level, the number of type I myofibers was significantly increased in the soleus and EDL muscles compared with *Rosa26*-MEK1 littermate controls (Fig. 3C-E).

**Figure 3.**
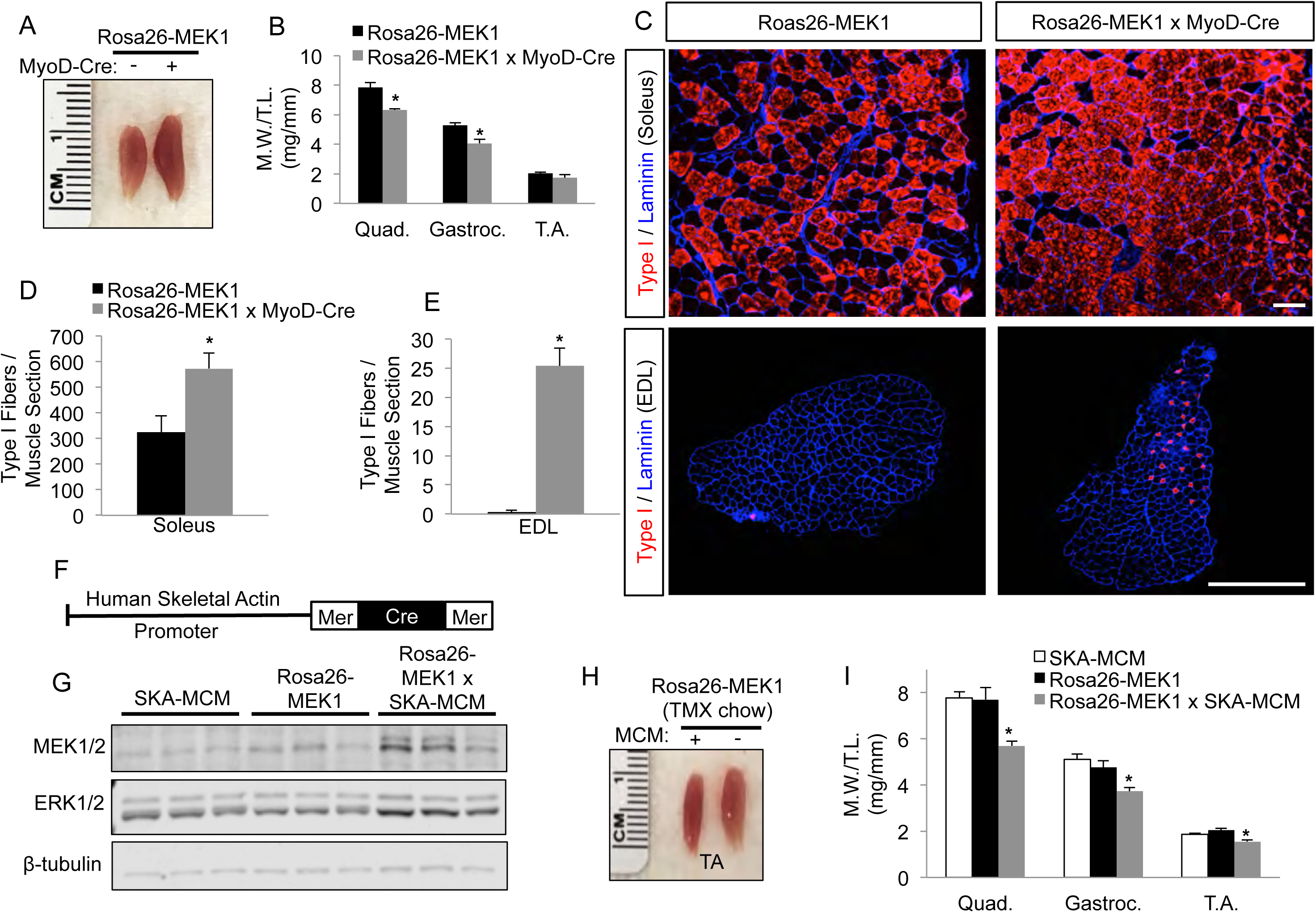
Expression of activated MEK1 in muscle progenitors or mature myofibers leads to a slow oxidative phenotype. (A) Representative image of the TA muscles from 6 month-old *Rosa26*-MEK1 animals with or without MyoD-Cre. (B) M.W./T.L. at 6 months of age for the Quad, Gastroc, and TA muscles from mice of indicated genotypes. n = 3-4 mice per group, ^*^P < 0.05. (C-E) Representative images from immunostained sections for type I fibers (red) and laminin (blue) with quantification from the soleus (C and D, n = 2-4 mice per group) and EDL (C and E, n = 3-5 mice per group) in the indicated genotypes of mice. Histological sections were cut from the mid-belly of the muscle to quantify total number of type I fibers per section. Scale bars: 500 μm. ^*^P < 0.05. (F) Schematic of the MerCreMer cDNA driven by the human α-skeletal actin promoter in generating muscle-specific transgenic mice. (G) Western blot analysis for total MEK1/2 and total ERK1/2 using lysate from the gastrocnemius muscle from mice of the indicated genotypes (n = 3). β-tubulin was used as a loading control. (H) Representative image of the TA muscle from 6 month-old *Rosa26*-MEK1 and *Roas26*-MEK1 × SKA-MCM animals. (I) M.W./T.L. at 6 months of age for the Quad, Gastroc, and TA muscles from *Rosa26*-MEK1 and *Rosa26*-MEK1 × SKA-MCM mice. n = 4-9 mice per group, ^*^P < 0.05 versus controls.

To determine whether a greater slow fiber phenotype could be induced in adult mice, we used a transgene in which the human skeletal α-actin promoter was used to drive the tamoxifen-regulated MerCreMer cDNA (SKA-MCM, Fig. 3F). Here we began administering tamoxifen to *Rosa26*-MEK1 × SKA-MCM mice as well as to control *Rosa26*-MEK1 littermates and SKA-MCM control mice at 2 months of age. Western blot analysis revealed an increase in total MEK1 and ERK1/2 proteins in Rosa26-MEK1 × SKA-MCM animals compared to controls (Fig. 3G). Even when activated MEK1 was induced in adult muscle, it still caused a slow-oxidative fiber-type switch and a reduction in muscle weights (Fig 3H,I). These results indicate that MEK1-ERK1/2 signaling induces a fast-to-slow myofiber transition in adult myofibers.

### Increased oxygen consumption in Rosa26-MEK1 MLC-Cre animals

Type I fibers are known to consume more oxygen, hence we performed indirect calorimetry in *Rosa26*-MEK1 × MLC-Cre mice to measure this metabolic indicator. Mice were housed in a sealed container with a motorized treadmill apparatus with an Oxymax system for gas analysis. Animals were acclimatized to the treadmill at the lowest speed setting (3m/min) for 10 min. During this initial period, we observed an increase in oxygen consumption in *Rosa26*-MEK1 × MLC-Cre mice compared to controls (Fig. 4A). Mice were then sprinted and we observed an even greater oxygen consumption differential between *Rosa26*-MEK1 × MLC-Cre mice compare with controls (Fig 4A). We also measured the respiratory exchange ratio (RER) to infer metabolic substrate utilization. RER values of 1 indicate preferential glucose oxidation, while RER values of 0.7 indicate preferential fatty acid oxidation. No differences in substrate utilization between *Rosa26*-MEK1 × MLC-Cre and control mice were observed during the exercise protocol (Fig. 4B). These results are consistent with a prominent shift in all musculature towards a slow-oxidative fiber phenotype with increased MEK1-ERK1/2 signaling.

**Figure 4.**
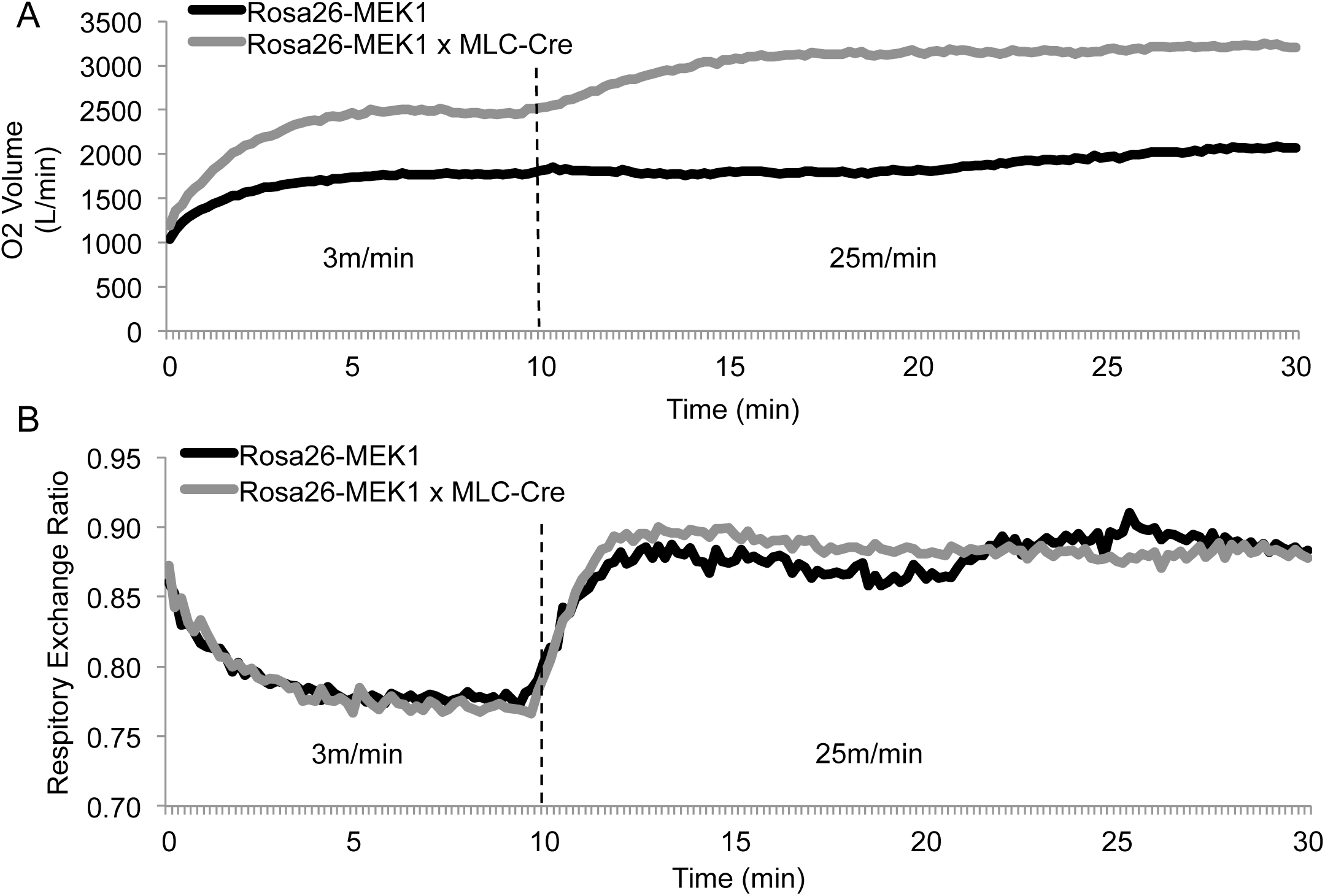
Increased oxygen consumption in *Rosa26*-MEK1 × MLC-Cre animals. (A) Oxygen consumption and (B) respiratory exchange ratio from 2 month-old *Rosa26*-MEK1 and *Rosa26*-MEK1 × MLC Cre mice during treadmill acclimatization (3 m/min) and exercise (25 m/min) period (n = 5).

### TA muscles from Rosa26-MEK1 × MLC-Cre mice are fatigue resistant

To evaluate how increased MEK1-ERK1/2 signaling impacted physiologic performance we evaluated muscle function using a whole leg immobilization preparation in 4 month-old *Rosa26*-MEK1 × MLC-Cre versus *Rosa26*-MEK1 control mice. The peak force produced by a single twitch contraction in the TA muscle was similar between both groups (Fig. 5A). However, we observed a 33% decrease in the absolute maximal peak tetanic force produced by the TA muscle from *Rosa26*-MEK1 × MLC-Cre mice compared to controls (Fig. 5B). Because the mild weight loss in all muscles from *Rosa26*-MEK1 × MLC-Cre mice we normalized the peak tetanic force to the cross-sectional area from the TA, which was still decreased by 18% versus control TA muscle (Fig 5C). These results are in agreement with those previously reported in other mouse models rich in type I fibers that have less maximal force generation capacity [25], again suggesting that increased MEK1-ERK1/2 signaling shifted muscles to a slow-oxidative program.

**Figure 5.**
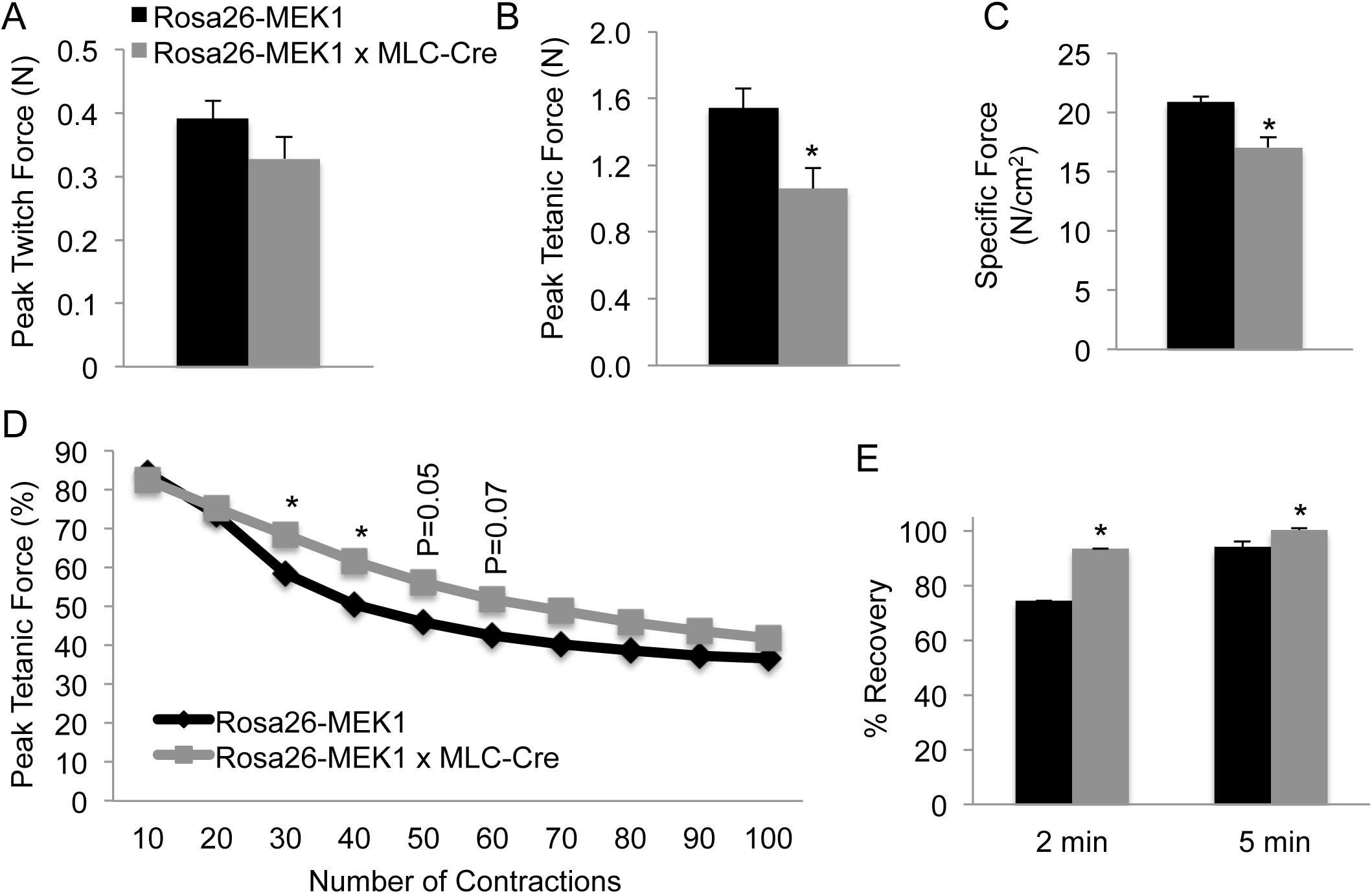
Skeletal muscles from *Rosa26*-MEK1 × MLC-Cre animals are fatigue resistance. (A) Peak twitch force and (B) peak tetanic force were measured from the TA muscle from 4 month-old *Rosa26*-MEK1 and *Rosa26*-MEK1 × MLC-Cre animals (n = 5, ^*^P < 0.05). (C) Normalized specific force measured from the TA muscle from mice of the indicated genotypes (n = 5, ^*^P < 0.05). (D) TA muscles from *Rosa26*-MEK1 and *Rosa26*-MEK1 × MLC-Cre were contracted 100 times to elicit fatigue. Peak tetanic forces are expressed as percentage of the pre-fatigue peak tetanic force. n = 4-5 mice per group, ^*^P < 0.05. (E) Peak tetanic force was recorded 2 min and 5 min post-fatigue to assess recovery in the indicated groups of mice. Data are presented as percentage of the pre-fatigue peak tetanic force. n = 3-4 mice per group, ^*^P < 0.05.

We next assessed if the increased proportion of type I fibers in muscles from *Rosa26*-MEK1 × MLC-Cre mice was associated with resistant to fatigue. *Rosa26*-MEK1 × MLC-Cre and *Rosa26*-MEK1 controls were subjected to a fatigue protocol in which we elicited 100 muscle contractions with only 3 s between each contraction. Whereas both groups showed a decrease in force production over time, the decrease in force from control *Rosa26*-MEK1 animals was more pronounced than in *Rosa26*-MEK1 × MLC-Cre animals (Fig. 5D). Following the fatigue protocol, we measured the maximal force produced by both groups at two different time points to assess muscle function recovery. Two minutes following the fatigue protocol, the TA muscle from *Rosa26*-MEK1 × MLC-Cre animals showed a 20% improvement in force recovery compared to *Rosa26*-MEK1 controls (Fig. 5E). At the 5 min post-fatigue time point, *Rosa26*-MEK1 × MLC-Cre animals had fully recovered and produced maximal forces identical to those observed prior to the fatigue protocol. In comparison, Rosa26-MEK1 control mice produced forces that were equal to 93% of their pre-fatigue maximal force, which was significantly less than *Rosa26*-MEK × MLC-Cre mice (Fig. 5E). These data demonstrate that increased MEK1-ERK1/2 signaling produces muscle that is more resistant to fatigue, indicative of a slow-oxidative switch.

## Discussion

Health benefits associated with endurance training are numerous; they include improved cardiovascular health [26], reduced obesity-linked insulin resistance [27] as well as improved cognitive function [28]. Increased capillary and mitochondrial content are two physiological adaptations associated with increased oxidative metabolism [1]. These adaptations can cause changes in metabolic enzyme profiles and expression changes in contractile protein isoforms. Ultimately, these adaptations to long-term endurance training lead to a fast-to-slow fiber-type transition. A proteomic screen revealed that some 562 proteins become phosphorylated in human skeletal muscle following exercise [29], suggesting that complex adaptive kinase-dependent signaling occurs in response to exercise. In the present study, we uncover a role for MAPK signaling in promoting a fast-to-slow fiber-type transition under basal conditions *in vivo.* The MEK1-ERK1/2 signaling pathway was shown to program a greater number of slow oxidative fibers in skeletal muscles, which now consume more oxygen and are more fatigue resistant compared with fast-glycolytic fibers.

ERK1 and ERK2 share 85% amino acid sequence identity, and each appear to phosphorylate the same substrates *in vitro* [30]. Despite their sequence similarity, the requirement for ERK1 versus ERK2 during development differs significantly. While MEK2 and ERK1 are dispensable for embryonic development, studies from gene-deleted mice revealed that MEK1 and ERK2 are essential for development, as loss of either gene product results in embryonic lethality [31]. We demonstrated here that MEK1-mediated activation of ERK2 but not ERK1, generated a pronounced slow-oxidative phenotypic switch in muscle. In our study, we used an antibody that detected an identical epitope present in ERK1 and ERK2, allowing us to conclude that ERK2 (lower band) is more abundantly expressed than ERK1 (upper band) in skeletal muscle, which we interpret as the likely mechanism for the differential effect on fiber-type switching downstream of MEK1 (Fig. 1B). This presumed effect is also consistent with a much more prominent upregulation of ERK2 in *Mapk3* null muscle versus ERK1 in the *Mapk1*-targeted mice (compare Fig 2A versus 2E). Our observations that MEK1-ERK1/2 signaling augments the slow-oxidative fiber-type program is actually opposite of the interpreted results of a previous report [17]. Shi et al transfected adult skeletal muscle with an MKP-1 expressing plasmid, whcih showed increased type IIa and type I fibers, indicating a more slow-oxidative program with ERK1/2 dephosphorylation [17]. However, MKP-1 is actually more specific for p38 MAPK and it can dephosphorylate all 3 major terminal MAPKs when overexpressed [18].

Type I fibers can be observed in muscles at birth that are traditionally fast-glycolytic, however these fibers are replaced by fast fibers as the muscles mature [32]. Neuromuscular junction development, increased mechanical load and thyroid hormone are all factors that contribute to a fiber-type switching during postnatal muscle maturation [1]. In our study, we have taken advantage of various muscle-specific Cre drivers to induce MEK1 activation during myogenesis and in adulthood. Regardless, of the temporal induction of MEK1-ERK1/2 signaling, we observed a shift towards a slow-oxidative program in all muscles examined in this study. We noted an increase in the number of type I fiber in the soleus muscle of *Rosa26*-MEK1 × MLC-Cre animals in spite of the fact that the MLC-Cre allele is more active in fast-twitch fibers compared with type I fibers. However, MyoD-Cre mouse also produced near identical results in promoting the slow fiber-type transition to Rosa26-MEK1 × MLC-Cre animals, and because the MyoD-Cre is active prior to developmental fiber-type specification is indicates that MEK1-ERK1/2 signaling acts independent of initial fiber-type identity.

Calcineurin is a calcium/calmodulin-regulated protein phosphatase that, once activated, can directly dephosphorylate members of the nuclear factor of activated T-cells (NFAT) transcription factor family [33]. Dephosphorylation of NFAT family members by calcineurin allows their translocation into the nucleus. Transgenic mice expressing an active form of calcineurin in skeletal muscle display a fast-to-slow fiber-type switch [34]. Conversely, loss of calcineurin A α/β isoforms leads to a down-regulation of the slow oxidative program in skeletal muscles [35]. Cross-talk between MEK1-ERK1/2 and the calcineurin-NFAT signaling pathways has previously been established in the heart where these pathways interact [36]. Immunoprecipitation experiments using protein lysate from cardiomyocytes revealed the formation of a protein complex involving MEK1-ERK2-calcineurin-NFAT. Sanna et al. further showed that NFAT binding activity was regulated by ERK1/2 *in vitro* [36]. However, we have not ascertained the downstream mechanism whereby increased ERK1/2 signaling leads to an induction of the slow fiber-type program, although the mechanism is likely combinatorial with many downstream effectors. We have shown that increased MEK1-ERK1/2 signaling is directly responsible for programming the fast-to-slow fiber-type program *in vivo* on many levels. This knowledge could be exploited for therapeutic or muscle performance advantage in going forward. Indeed, augmenting the slow-oxidative fiber-type program would provide a metabolic benefit, provide greater endurance performance and likely also benefit patients with muscular dystrophy [25, 37, 38].

## Materials and Methods

### Animal Models

Animal experiments performed in the study were approved by the Institutional Animal Care and Use Committee of the Cincinnati Children’s Hospital Medical Center. Mouse lines used in the study were: *Mapk3*^-/-^ (ERK1) mice [39], *Mapk1^f/f^* (ERK2) mice [40] and mice harboring a Cre-dependent constitutively active MEK1 cDNA inserted into the *Rosa26* locus [41] (Jax Stock No: 012352). *Mapk1^f/f^* animals and those with the constitutively active MEK1 allele were crossbred with mice expressing Cre recombinase under the control of the myosin light chain 1f (*Myl1*, referred to as MLC-Cre throughout) genomic locus (knock-in) [21]. Additionally, we crossed the mice with the constitutively active MEK1 cDNA with either MyoD^iCre^ gene-targeted animals [24] or with SKA-MCM transgenic mice, which were generated by subcloning a cDNA coding for a Cre recombinase protein fused with two mutant estrogen-receptor ligand-binding domains on either side [42] into the pcDNA3.1 vector driven by the human α-skeletal actin promoter [43]. Tamoxifen was administered to *Rosa26*-MEK1 × SKA-MCM and *Rosa26*-MEK1 controls via intraperitoneal injections at 2 months of age for 5 consecutive days (75 mg/kg) using pharmaceutical-grade tamoxifen (Sigma) dissolved in corn oil (Sigma). Subsequently, the mice were fed a diet containing 400 mg/kg tamoxifen citrate (Envigo, TD.55125) for the duration of the study. Mice were sacrificed by isoflurane inhalation followed by cervical dislocation. Skeletal muscles were either embedded in tragacanth (Sigma), frozen in liquid-nitrogen cooled 2-methylbutane for downstream histological analyses or immediately flash frozen and stored at −80°C for biochemical analyses.

### Immunofluorescence

Histological cross-sections (8 μm) were collected from skeletal muscles using a cryostat and maintained in blocking solution (10% goat serum diluted in phosphate buffered saline (PBS)) for 30 min in a humid chamber. Slides were stained overnight at 4°C with PBS containing anti-MYH7 (BA-F8, 1:50, Developmental Studies Hybridoma Bank) primary antibody to detect type I fibers and with anti-laminin antibody (1:200, Sigma) to delineate myofiber outlines present in a muscle section. Primary antibodies were visualized using Alexa-568 goat anti-mouse IgG2b (Invitrogen) and Alexa-405 goat anti-rabbit IgG secondary antibodies diluted 1:500 in PBS. Immunofluorescence images were captured using a Nikon Eclipse Ti microscope. Entire muscle sections were analyzed for type I positive fibers. All experiments were performed in a blinded fashion whereby the experimenter was only made aware of the genotypes after the quantification was performed.

### Western Blotting

Muscles were homogenized and lysates prepared as previously described [44]. Proteins were resolved by SDS-PAGE, transferred to Immobilon-FL membranes (Millipore) and incubated overnight with antibodies against ERK1/2 (1:2000, Cell Signaling Technology), MEK1/2 (Cell Signaling Technology), anti-MYH7 (1:50), GAPDH (1:500 000, Fitzgerald) and β-tubulin (E7, 1:200, Developmental Studies Hybridoma Bank). Membranes were then incubated with IRDye secondary antibodies (1:6000, LI-COR Biosciences) and visualized using an Odyssey CLx imaging system (LI-COR Biosciences).

### Measurement of basal metabolic parameters

To monitor oxygen consumption during exercise, a sealed motorized treadmill was used. The treadmill had adjustable speed and inclination settings and was equipped with an electric shock-delivering grid. During the acclimatization period, the treadmill was set at its slowest speed (3 m/min) for 10 min followed by an exercise period (25 m/min) lasting 20 min. Electric shock intensity was set to 1 mA and the inclination was set to 5%. Fresh air was delivered with an electric pump and experiments were performed at 23°C. Gas samples from the treadmill chamber were collected every 15 s and analyzed by the Oxymax system (Columbus Instruments) for measurement of VO_2_ and VCO_2_. Respiratory exchange ratio (RER) was calculated as VO_2_/VCO_2_ with the CLAMS data examination tool (Columbus Instruments).

### In situ TA isometric force measurement

Mice were anesthetized with isofluorane and a 1.5 cm incision was made along the femur exposing the sciatic nerve that allowed for the identification of the tibia branch of this nerve. Both the tibia branch and the sciatic nerve were cut, leaving the peroneal branch of the nerve intact for stimulation. The skin running from the ankle to the thigh was cut and removed to expose the TA muscle. The mouse was then placed in a supine position on the muscle testing apparatus while a radiant lamp was used to maintain the animal’s body temperature. The knee was immobilized at the limb clamp (Aurora Scientific) and the foot was fixed using surgical tape. The lower third of the TA muscle was dissected off the tibia and the tendon was resected and attached to the lever arm using silk suture to measure force production (Aurora Scientific, 305C). The sciatic nerve was then placed on two needle electrodes and stimulated to determine the maximal isometric twitch force with 0.2 ms pulses at 50 mA. The muscle was stretched until the optimal length was reached that allowed for maximal force production. Next, the maximal isometric tetanic force was measured by increasing the frequency from 25 to 200 Hz. The duration of the contractions was set at 350 ms and a contraction was elicited every 2 min.

Muscle fatigue was achieved by increasing the contraction rate to one contraction every 3s for 100 contractions. The maximal tetanic force was measured after each contraction during the fatigue protocol. In addition, we measured the maximal tetanic force at 2 and 5 min following the fatigue protocol to assess post-fatigue recovery. Muscle weight and length were measured to calculate the physiological cross-sectional area using the following formula: [*muscle mass (g)/1.06 (fiber density) × muscle length (cm) × 0.6 (muscle length to fiber length ratio)*]. To normalize the maximal force production, we calculated the specific force by dividing the absolute tetanic force by the physiological cross-sectional area of the TA muscle.

### Statistics

A one-way analysis of variance was used to determine if there was a significant difference in experiments with more than 2 groups; a Tukey’s post hoc test was performed to compare individual groups (Prism Software). Significant differences between 2 groups were determined using an unpaired Student’s *t* test (Stats Plus Software). Significance was set at P<0.05 for all experiments. All results are presented as mean plus the standard error of the mean.

## Acknowledgements

J.D.M. was funded by grants from the National Heart Lung and Blood Institute of the National Institutes of Health, and by the Howard Hughes Medical Institute

