## Supplemental figures for "Mitogen-Activated Protein Kinase-Dependent Fiber-Type Regulation in Skeletal Muscle"

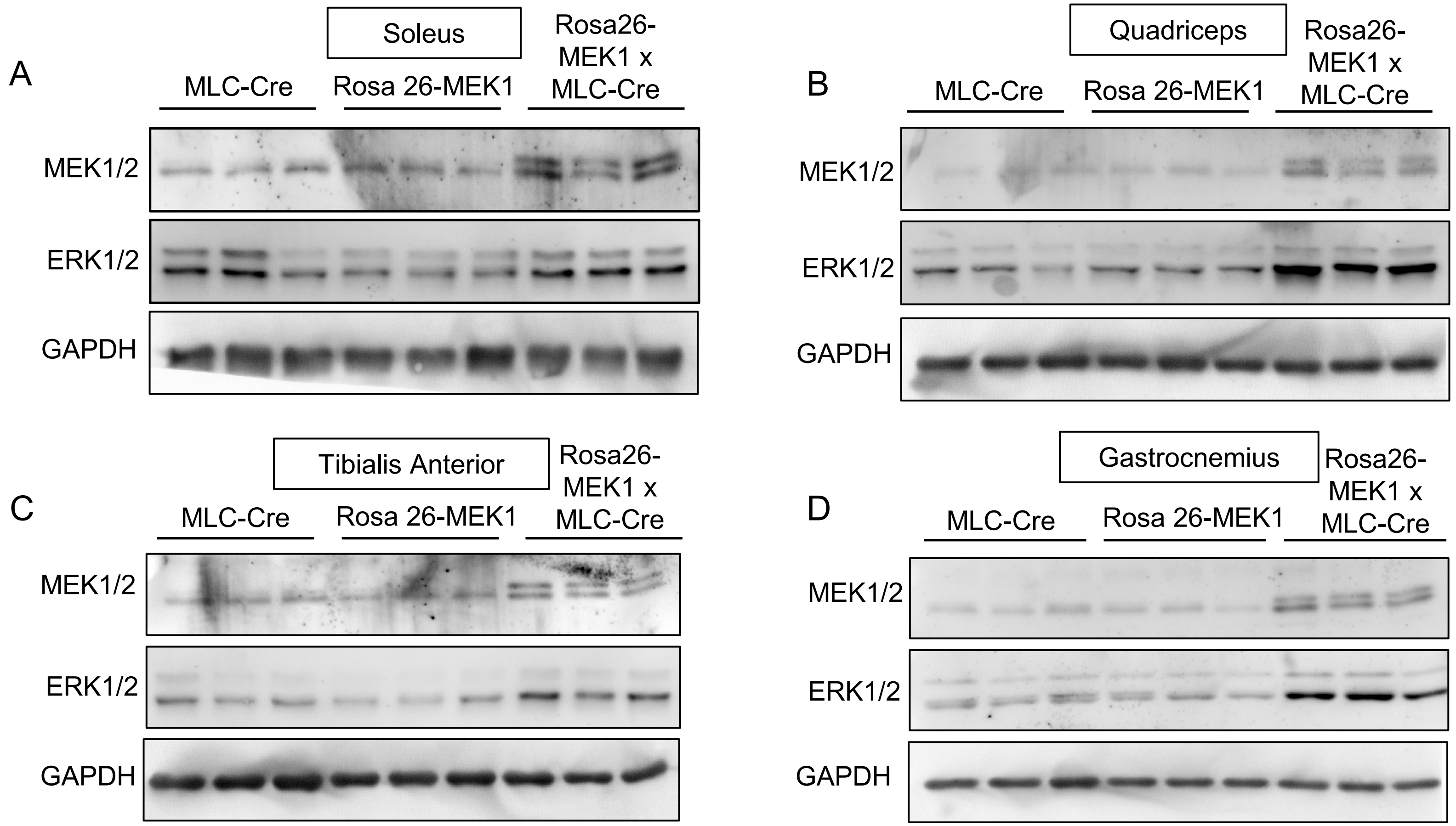


**Supplementary Figure 1. Activation of ERK1/2 in skeletal muscle.**
Western blot analysis for total MEK1/2 and total ERK1/2 using muscle lysate from (A) the soleus muscle, (B) the quadriceps muscle, (C) the tibialis anterior muscle and (D) in the gastrocnemius muscle (n = 3). GAPDH was used a loading control.

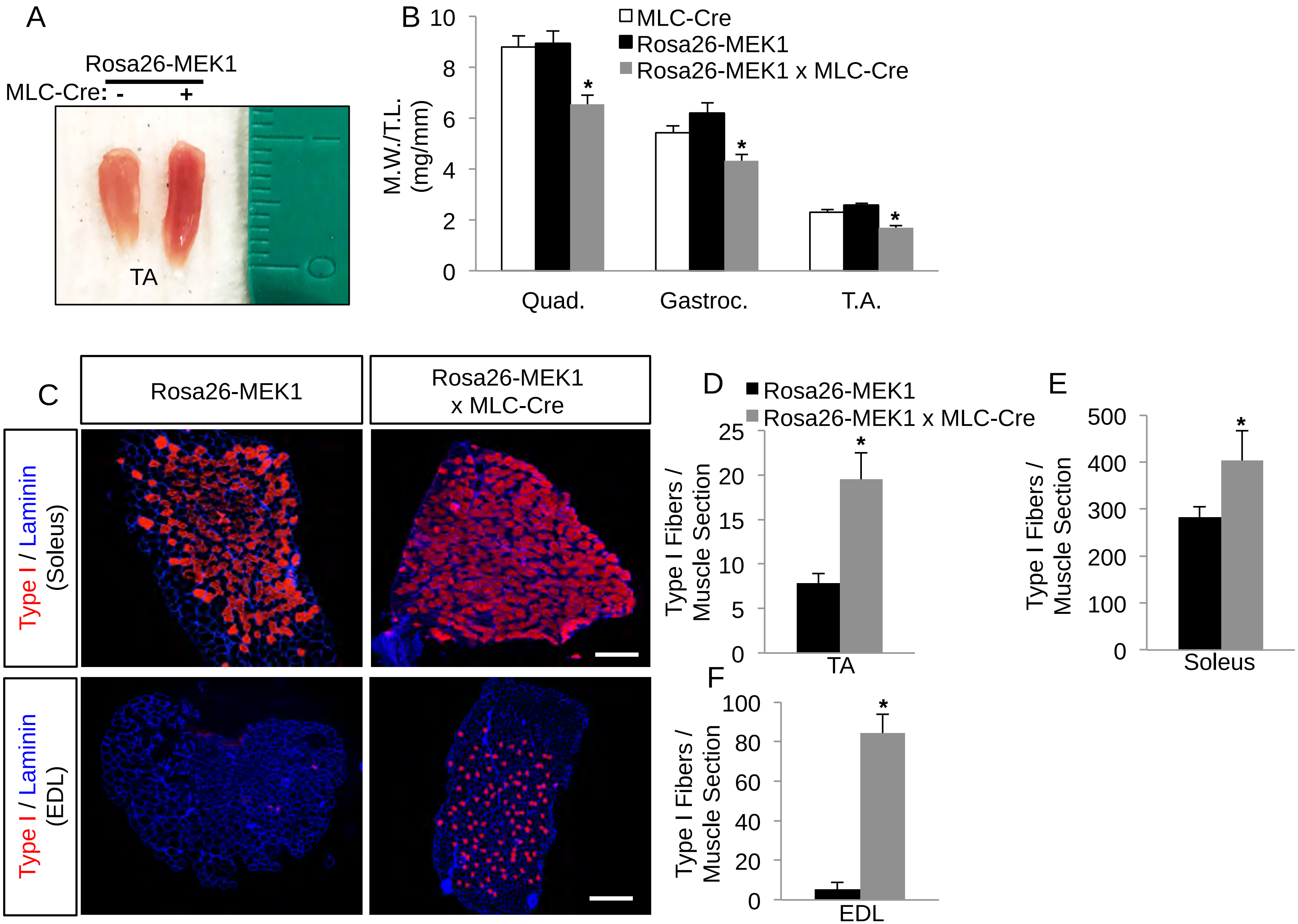


**Supplementary Figure 2. MEK1-ERK1/2 signaling increases the number of type I fibers.**
(A) Representative image of the tibialis anterior (TA) muscles from 2 month-old *Rosa26*-MEK1 and *Rosa26*-MEK1 x MLC-Cre animals. (B) Muscle-weight normalized to tibia-length (M.W./T.L.) at 2 months of age from mice of the indicated genotypes. Muscles analyzed are shown. n = 5-10 mice per group. *P < 0.05 versus controls. (C) Representative immunohistochemical stained images from the soleus and extensor digitorum longus (EDL) muscles showing MHY7 (red) positive type I fibers and laminin expression (blue) used to identify all myofibers present in sections from mice of the indicated genotypes. Scale bars: 500 µm. (D-F) Quantitation of type I fibers in sections from the TA muscle (D), the soleus muscle (E) and the EDL muscle (F) of *Rosa26*-MEK1 and *Rosa26*-MEK1 x MLC-Cre animals at 2 months of age. n = 4-6 mice per group. *P < 0.05.
